# Measurements of translation initiation from all 64 codons in *E. coli*

**DOI:** 10.1101/063800

**Authors:** Ariel Hecht, Jeff Glasgow, Paul R. Jaschke, Lukmaan Bawazer, Matthew S. Munson, Jennifer Cochran, Drew Endy, Marc Salit

**Affiliations:** Joint Initiative for Metrology in Biology, Stanford, CA; Genome-scale Measurements Group, National Institute of Standards and Technology, Stanford, CA; Department of Bioengineering, Stanford, CA; Department of Chemistry and Biomolecular Sciences, Macquarie University, Sydney, NSW, Australia

**Author notes:** To whom correspondence should be addressed. Correspondence may also be addressed to Drew Endy. These authors contributed equally (Ariel Hecht, Jeff Glasgow, Paul R. Jaschke, Lukmaan Bawazer).

## Abstract

Our understanding of translation is one cornerstone of molecular biology that underpins our capacity to engineer living matter. The canonical start codon (AUG) and a few near-cognates (GUG, UUG) are typically considered as the “start codons” for translation initiation in *Escherichia coli* (*E. coli*). Translation is typically not thought to initiate from the 61 remaining codons. Here, we systematically quantified translation initiation in *E. coli* from all 64 triplet codons. We detected protein synthesis above background initiating from at least 46 codons. Translation initiated from these non-canonical start codons at levels ranging from 0.01% to 2% relative to AUG. Translation initiation from non-canonical start codons may contribute to the synthesis of peptides in both natural and synthetic biological systems

## INTRODUCTION

The translation of messenger RNA (mRNA) to protein is one of the fundamental processes in biology. Elucidation of translation represents a central aim of molecular biology. Control of translation is critical to enable precision bioengineering. One key regulator of translation is the initiation or “start” codon, which is the three mRNA nucleotides that bind to *N*-formylmethionyl transfer RNA (tRNA^fMet^) (1, 2). The most common bacterial start codons are AUG, GUG and UUG (2, 3), initiating translation of 83%, 14%, and 3%, respectively, of known *E. coli* genes (4). Similar percentages can be found throughout the bacterial domain (5).

The occurrence of non-canonical start codons (defined here as any codon other than AUG, GUG and UUG) in known genes is very rare. Only two non-canonical start codons have been confirmed in *E. coli: infC* (6) and *pcnB* (7) both begin with AUU. While there are similarly small numbers of confirmed non-canonical start codons in eukaryotes (8), recent ribosomal footprinting studies with yeast and mammalian cells suggest that non-canonical translation initiation may be much more prevalent than previously thought (9–12).

Translation initiation is an intricate process that has been detailed in many excellent reviews (e.g., 13–18). Of the mRNA sequences that regulate translation initiation, the impact of variation within the 5’ untranslated region (5’ UTR) (19) and the Shine-Dalgarno (SD) sequence [also known as the ribosome binding site (RBS)] (20) have been systematically quantified (13, 21–24). However, the start codon itself has not yet been systematically explored.

Codons can adopt a number of different roles in translation. As examples: organisms across all domains of life naturally reassign one of the three canonical stop codons (UAA, UAG and UGA) to code for amino acids (25); in wild microbes 13 different codons have evolved to code for the 21st proteinogenic amino acid selenocysteine (26); and, in engineered *E. coli,* 58 of the 64 codons were successfully reassigned to code for selenocysteine (27), and the UAG stop codon was removed from the genome to allow its reassignment to unnatural amino acids (28).

Ongoing improvements in DNA synthesis (29), sequencing (30) and assembly (31, 32), and the creation of a variety of bright fluorescent proteins (33), have enabled a wave of recent efforts to systematically explore, modify, and fine-tune genetic regulatory elements (34–38). These efforts have led to, for example, new designs for promoters (39–41), repressors (42, 43), ribosome binding sites (39, 44, 45), insulators (46), and terminators (47). Systematic explorations of the regions immediately upstream (22, 23) and downstream (48–51) of the start codon in *E. coli* have revealed significant impacts of the sequence surrounding the start codon on translation efficiency. However, only a few studies have explored initiating translation from the near-cognates of AUG (52, 53), and none appear to have explored initiating translation from all 64 codons.

## MATERIALS AND METHODS

### Bacterial culture

All strains were grown in lysogeny broth (LB) or on LB agar, supplemented with 50 μg mL^−1^ kanamycin, 100 μg mL^−1^ carbenicillin, 100 μg mL^−1^ ampicillin, or 25 μg mL^−1^ chloramphenicol (Sigma) for selection. ^i^ Plasmids were isolated using a “QIAQuick Miniprep” kit (Qiagen). PCR reaction products were purified using a GeneJet Gel extraction kit (Thermo Scientific) or NucleoSpin Gel and PCR Clean-Up kit (ClonTech). Plasmids and PCR products were sequenced using Sanger sequencing (Elim Biopharma or MCLab). All PCR and cloning reactions were performed on a S1000 Thermal Cycler (Bio-Rad).

### Construction of T7-sfGFP plasmids

A library of 64 plasmids was created where the start codon (ATG) in the superfolder GFP (sfGFP) coding sequence was replaced with each of the 64 codons. A pET20b vector with a pBR322 origin of replication and an ampicillin-resistance cassette was used as the plasmid backbone (Supplementary Table S2). The sfGFP transcript had a strong ribosome binding site (AGGAGA), and the spacer between the RBS and the start codon (TAAATAC) was designed to prevent the creation of out-of-frame canonical start codons and achieve optimal RBS-start codon spacing (54). 31 variants of the 64-member library were created by one-pot golden gate cloning, and the remaining 33 were created in parallel reactions via plasmid amplification followed by blunt-end ligation. Bacterial cultures for plasmid construction were grown in LB supplemented with ampicillin. Additional details about the cloning methods used can be found in the Supplementary Information.

### Construction of RhaP_BAD_-sfGFP and RhaP_BAD_-NanoLuc plasmids

A set of 12 codons (AUG, GUG, UUG, AUA, AUC, AUU, CUG, CAU, CGC, GGA, UAG and UGC) were selected for further exploration as potential start codons in three different expression cassettes: sfGFP under control of the native *E. coli RhaP_BAD_* rhamnose-inducible promoter on a p15A plasmid, NanoLuc luciferase (Promega) under control of *RhaP_BAD_* on a p15A plasmid, and NanoLuc luciferase under control of *RhaP_BAD_* on a single-copy bacterial artificial chromosome (BAC) (55). Sequence information (Supplementary Table S2) and additional details about the cloning are available in the Supplementary Information.

### Construction of RhaP_BAD_-Luciferase BACs

To better mimic physiologically-relevant expression conditions, we cloned the NanoLuc luciferase expression cassettes into a bacterial artificial chromosome (BAC), which constitutively expresses a single-copy oriS origin of replication (55), with chloramphenicol and ampicillin resistance markers. The BAC carried a second origin of replication (oriV), which was high-copy, and inducible with 1% arabinose. We used oriV for improving yields of BAC isolation and cloning reactions. NanoLuc expression cassettes with each of the 12 different start codons were amplified from the p15A vectors that were constructed in the previous step. Additional details about the cloning methods used can be found in the Supplementary Information.

### Culture growth conditions for assay measurements

LB agar plates with the appropriate antibiotics were streaked from frozen glycerol stocks and incubated overnight at 37 °C. Plates were stored at 4 °C until ready for use. Plates were discarded after two weeks of storage at 4 °C. Three individual colonies for each construct were used to inoculate 300 μL of LB containing the appropriate antibiotics: carbinecillin and chloramphenicol (pET20b-sfGFP vectors in BL21(DE3)pLysS cells), or kanamycin (all others) in a 96-well deep well culture plate (VWR). The plate was sealed with an AeraSeal gas-permeable microplate seal (E&K Scientific) and grown overnight at 37 °C in a Kuhner LT-X (Lab-Therm) incubator shaking at 460 rpm with 80% relative humidity.

Negative control strains were prepared for each experiment to measure background signal. For experiments measuring sfGFP expression, an empty vector backbone with no expression cassette (pET20b) or a plasmid expressing a non-fluorescent silicatein gene (p15A) was used. This control plasmid was transformed into the same strains as used for measurement for use as a negative control strain. For experiments measuring NanoLuc luciferase expression, the same strains used for measurement with a vector expressing sfGFP (p15A or BAC) were used as a negative control.

### Fluorescence measurements

After the overnight growth, cultures were sub-cultured 1:100 into 300 μL of EZ Rich Defined Medium (Teknova) with the same antibiotics as the initial culture and grown for 2 hours under the same conditions. Expression of sfGFP was then induced by supplementing the cultures with 5 mmol/L inducer (IPTG for the pET20b-sfGFP constructs or rhamnose for the *RhaP_BAD_*-sfGFP constructs), and the cells were grown for an additional 5 hours (pET20b-sfGFP constructs) or 13 hours (pRha-sfGFP constructs). For plate reader measurements, 50 μL of each culture were transferred to a CELLSTAR black, clear-bottom 96-well plate (Grenier, #M0562). 150 μL of 1X PBS (Fisher Scientific) was added to each well. Cultures were analyzed on a BioTek Synergy H4 plate reader. Absorbance at 600 nm (OD_600_) was measured to estimate culture density, followed by fluorescence (ex. 485 nm/em. 510 nm, bw = 9.0 nm, sensitivity = 65).

For flow cytometry, cells were diluted 1/100 in phosphate buffered saline (20mmol/L phosphate, 0.85%NaCl) and measured on a BD LSRII instrument using a FITC fluorescence channel (488 nm excitation laser with a 525/50 emission band-pass filter). Detector voltages through forward scattering, side scattering, and FITC channels were set using white cells as a negative control and pET-sfGFP(ATG) clones as a positive control. Measured events were triggered on a side-scattering threshold and 30,000 events were measured from each cell culture. Two measurement sets were acquired, at high-and low-gain through the FITC channel. At high gain, the mean sfGFP signal from three highest expressing samples (ATG, GTG, TTG) were set off-scale, above the upper detection limit such that the remaining samples were within the range of detection. At low gain, the mean signals from highest expressing samples were within the measurable range, but means for the lowest expressing samples were off-scale below the lower detection limit. Flow cytometry data were processed in R (56), using the flowCore package (57) to remove negative near-baseline (noise) values and log transform the data, and ggplot2 (58) to generate violin plots.

### Luminescence measurements

After the overnight growth, cultures were subcultured 1:100 into 300 μL of LB with kanamycin and grown for 2 hours under the same conditions. Expression of NanoLuc was induced by supplementing the cultures with 4 mmol/L rhamnose, and then the cultures were incubated overnight as previously described. After the second overnight growth, 50 μL of each culture were transferred to a CELLSTAR black, clear-bottom 96-well plate. 150 μL of 1X PBS (Fisher Scientific) was added to each well. OD_600_ was measured as described above.

NanoLuc luciferase luminescence measurements were performed using the Nano-Glo Luciferase Assay System kit (Promega, #N1110), following the manufacturer’s protocol. Lysis buffer was prepared by dissolving 37.5 mg of Egg White Lysozyme (Sigma-Aldrich) in 7.5 mL of 1X Glo-Lysis Buffer (Promega). 20 μL of cell culture was diluted in 180 μL of lysis buffer and incubated at room temperature for 10 minutes. Assay buffer was prepared by mixing 9.8 mL of NanoLuc Assay Buffer (Promega) with 200 μL of NanoLuc Assay Reagent (Promega). 5 μL of lysed cells were mixed with 195 μL of assay buffer on a CELLSTAR white, opaque-bottom 96-well plate and incubated for 5 minutes at room temperature. Luminescence was measured on a BioTek Synergy H4 plate reader for all visible wavelengths for 1 second at a gain of 100. To minimize carry-over signal from adjacent wells, codons were separated by at least one row and known high-expressing codons (AUG, GUG, and UUG) were read separately.

### Data Analysis

Raw fluorescence and optical density measurements for each cell culture were imported into R. Wells filled only with EZ RDM were used to subtract background optical density, and negative control cells were used to subtract background fluorescence or luminescence. Per-cell fluorescence or luminescence was calculated by dividing the background-subtracted fluorescence by the background-subtracted optical density. Mean per-cell fluorescence or luminesce for each start codon in each expression strain was calculated by averaging the per-cell measurements from three biological replicates. For each expression system, per-cell expression was normalized by the per-cell expression from the AUG start codon to facilitate comparison of relative expression from different expression systems. The significance of the translation initiated from each codon was determined by comparing the expression measured from all of the codons in each experiment to the negative control using Dunnett's test (59), a statistical method for comparing multiple treatments to a single control, using the R multcomp package (60).

### Mass Spectrometry to confirm sfGFP

#### GFP expression and purification

Five GFP start codon variants were selected from different expression levels (ATC, ACG, CAT, GGA, and CGC). Genes for these were recloned into pET20b with a C-terminal 6x His-tag. These plasmids were transformed into BL21(DE3) pLysS and expressed on a larger scale. A single colony was used to inoculate 3 ml of LB supplemented with ampicillin and chloramphenicol and shaken at 37 °C overnight. The overnight culture was used to inoculate 30 mL of the same media 1:100 and grown for 2 hours at 37 °C, followed by induction with 1 mmol/L IPTG (final). The cells were allowed to express the protein at 37 °C for 5 hours and harvested by centrifugation for 10 minutes at 4000 x g. The cells were resuspended and lysed in 1 mL BPER with 0.6 mg/mL lysozyme, vortexed, and incubated for 10 minutes at room temperature. DNase (1 unit) was added and further incubated with frequent vortexing for 10 minutes. The lysate was then centrifuged at 4 °C for 10 minutes at 15,000 x g. The clarified lysate was run over 150 μL of nickel resin preequilibrated with 2xPBS (24 mmol/L sodium phosphate buffer, 274 mmol/L NaCl, 54 mmol/L KCl) with 20 mmol/L imidazole at pH 7.5. The column was washed with 4 mL of 2xPBS with 20 mmol/L imidazole. Green protein was eluted with 400 μl of the same buffer with 300 mmol/L imidazole.

#### Sample preparation for LC-MS

50 μL volume of each of the protein solution samples was aliquoted, followed by protein precipitation with the addition of 250 μL -80 °C acetone. Samples were stored at -80 °C for 1 hr, followed by light vortex and spun at 12,500 rpm for 12 minutes at 4 °C. The supernatant was decanted and the protein pellet was le ft to dry under the chemical hood for 20 minutes at room temperature. The protein pellet was re-suspended in 65 μL 50mmol/L ammonium bicarbonate 0.2% protease max (Promega) followed by vortex and sonication until the pellet was fully suspended. DTT was added to a final concentration of 5mmol/L, incubated on a heat block at 55°C for 30 minutes followed by alkylation with the addition of propionamide to a final concentration of 10 mmol/L for 30 minutes at room temperature. 2 μg of Asp N (Promega) was reconstituted in the vendor specific buffer and 0.5 μg added to each sample, followed by overnight digestion at 37°C. Formic acid was added to 1% and the acidified digest was C18 stage tip purified (Nest Group) using microspin columns and dried in a speed vac.

#### LC-MS

Peptide pools were reconstituted and injected onto a C18 reversed phase analytical column, 10 cm in length (New Objective). The UPLC was a Waters NanoAcquity, operated at 600nL/min using a linear gradient from 4% mobile phase B to 35% B. Mobile phase A consisted of 0.1% formic acid, water, Mobile phase B was 0.1% formic acid, water. The mass spectrometer was an Orbitrap Elite set to acquire data in a high/high data dependent fashion selecting and fragmenting the 10 most intense precursor ions in the HCD cell where the exclusion window was set at 60 seconds and multiple charge states of the same ion were allowed.

#### LC-MS data analysis

MS/MS data were analyzed using Preview and Byonic v2.6.49 (ProteinMetrics). Data were first analyzed in Preview to verify calibration criteria and identify likely post-translational modifications prior to Byonic analysis. All analyses used a custom. fasta file containing the target protein sequence, and were searched with a reverse-decoy strategy at a 1% false discovery rate. Byonic searches were performed using 12 ppm mass tolerances for precursor and fragment ions, allowing for semi-specific N-ragged tryptic digestion. The resulting identified peptide spectral matches were then exported for further analysis.

### Bioinformatics analysis

To analyze initiation codon annotations from model bacterial species, 79 complete genome sequences were collected from the National Center for Biotechnology Information databases (Supplementary Table S3). Initiation codon sequences were extracted from annotated features and compiled into a comprehensive list of 85,119 entries with Accession Number, Start Codon, Locus Tag, Gene Name, and Gene Product Name extracted from GenBank annotation features ID, sequence, qualifier(locus_tag), qualifier(gene), and qualifier(product) respectively. After removal of entries due to pseudogenes and misannotations a set of 84,897 entries remained (Supplementary Table S5) for analysis of initiation codon frequencies across the replicons of model bacterial species.

RNA folding simulations of transcripts from measured plasmids was performed using both NUPACK (61) and KineFold (62). NUPACK was run using default parameters. KineFold was run using default parameters except 3 ms co-transcriptional folding parameter for pET20b simulations and 20 ms co-transcriptional folding parameter for p15A and BAC simulations to account for the different RNA polymerases transcribing these vectors *in vivo.*

## RESULTS

We were first motivated to explore non-canonical start codons when we attempted to silence translation of a dihydrofolate reductase (DHFR) gene. We changed the start codon from AUG to UUG, GUG, AUA, or ACG. Surprisingly, we detected significant DHFR expression in recombinant bacterial extract (63) from all five codons (Supplementary Figure S1). We initially wondered whether our results were an artifact of *in vitro* translation; similar results reported in rabbit reticulocyte lysate suggested that our observations might merely be due to *in vitro* translation (64).

We analyzed 34 well-annotated bacterial genomes (65) to determine which of the 64 codons have been annotated as start codons (Supplementary Table S3). Our approach was similar to previous efforts (5) but with a focus on well-annotated genomes. Our analysis indicated that the vast majority of annotated open reading frames have ATG (81.8%), GTG (13.8%), or less frequently, TTG (4.35%) as the start codon, although CTG, ATT, ATC, and ATA are also annotated as start codons (Table 1 and Supplementary Table S5).

**Table 1.**
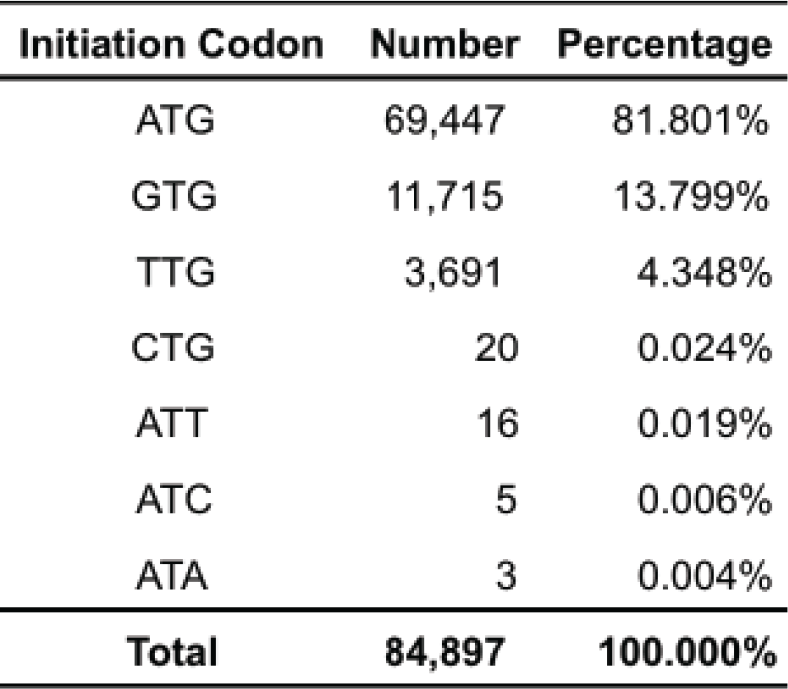
Annotated Initiation Codons in Model Bacterial Genomes. Start codons extracted from annotated features of 79 bacterial genome and plasmid sequences.

| **Initiation Codon** | **Number** | **Percentage** |
| --- | --- | --- |
| ATG | 69,447 | 81.801% |
| GTG | 11,715 | 13.799% |
| TTG | 3,691 | 4.348% |
| CTG | 20 | 0.024% |
| ATT | 16 | 0.019% |
| ATC | 5 | 0.006% |
| ATA | 3 | 0.004% |
| **Total** | **84,897** | **100.000%** |

We designed a set of four plasmids with different copy numbers, promoters, and reporters to experimentally quantify translation initiated from all 64 codons in *E. coli* (Figure 1). First, we measured the translation of superfolder GFP (sfGFP) initiated from all 64 codons. Expression was driven by a T7 promoter and a strong ribosome binding site (AGGAGA) on a high-copy pET20b vector (Figure 1A). The spacer sequence between the RBS and the start codon (UAAAUAC) was designed to be the optimal length for promoting translation initiation (54), and also to prevent the inadvertent creation of an in-frame or out-of-frame canonical start codon.

**Figure 1.**
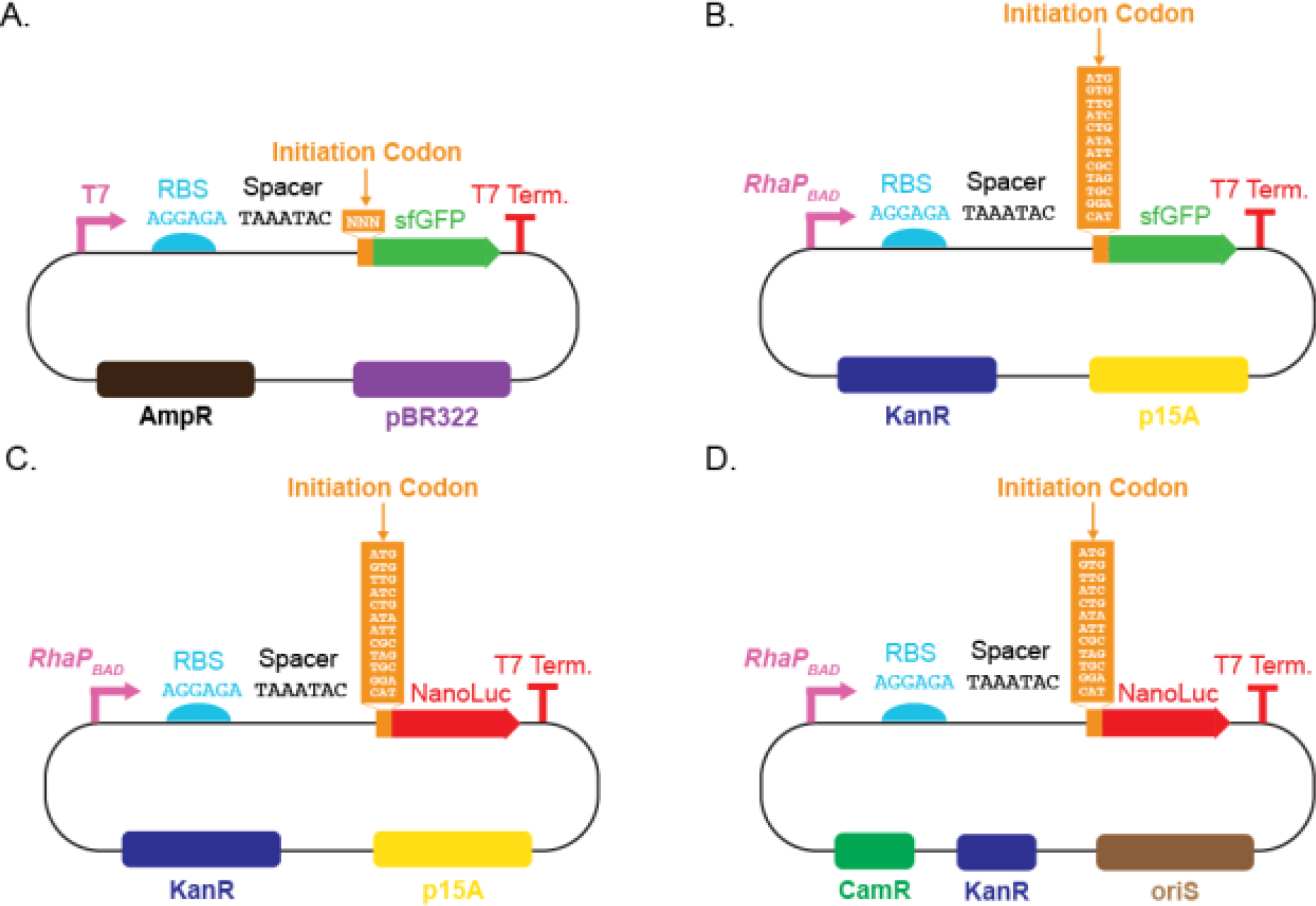
Plasmid sets used to measure translation initiation from non-canonical start codons. Plasmids varied in origin of replication (copy number), promoter, and reporter gene characteristics. (A) Set of 64 pET20b derived plasmids containing medium-copy pBR322 origin, T7 promoter, and superfolder GFP reporter. (B) Set of 12 plasmids containing low-copy p15A origin, *RhaP_BAD_* rhamnose-inducible native *E. coli* promoter, and sfGFP reporter. (C) Set of 12 plasmids containing low-copy p15A origin, *RhaP_BAD_* rhamnose-inducible native *E. coli* promoter, and NanoLuc^®^ luciferase reporter. (D) Set of 12 plasmids containing single-copy *oriS* bacterial artificial chromosome (BAC) origin of replication, *RhaP_BAD_* rhamnose-inducible native *E. coli* promoter, and NanoLuc^®^ luciferase reporter. Full construct sequences are available (Supplementary Table S2).

We measured fluorescence and absorbance via a plate reader. A strain carrying a plasmid without the sfGFP gene was used as a negative control. We calculated the mean per-cell fluorescence (fluorescence divided by OD_600_) for strains expressing sfGFP with each of the 64 codons inserted in place of the start codon in the sfGFP coding sequence. We normalized the mean per-cell fluorescence measured from each culture by the mean per-cell fluorescence measured from the culture expressing sfGFP with the canonical AUG start codon (Figure 2). The expression of sfGFP initiated from each culture was compared to the expression of the negative control using Dunnett's test, a method for comparing multiple treatments to a single control (59), assuming equal variance. Expression initiated from a codon was considered to be significant if the adjusted *p* value was less than 0.05 (red asterisks, Figure 2). Of the 64 start codons tested, translation initiated from 46 at a level significantly greater than the negative control. There was no significant correlation between fluorescence and cell abundance (Supplementary Figure S2).

**Figure 2.**
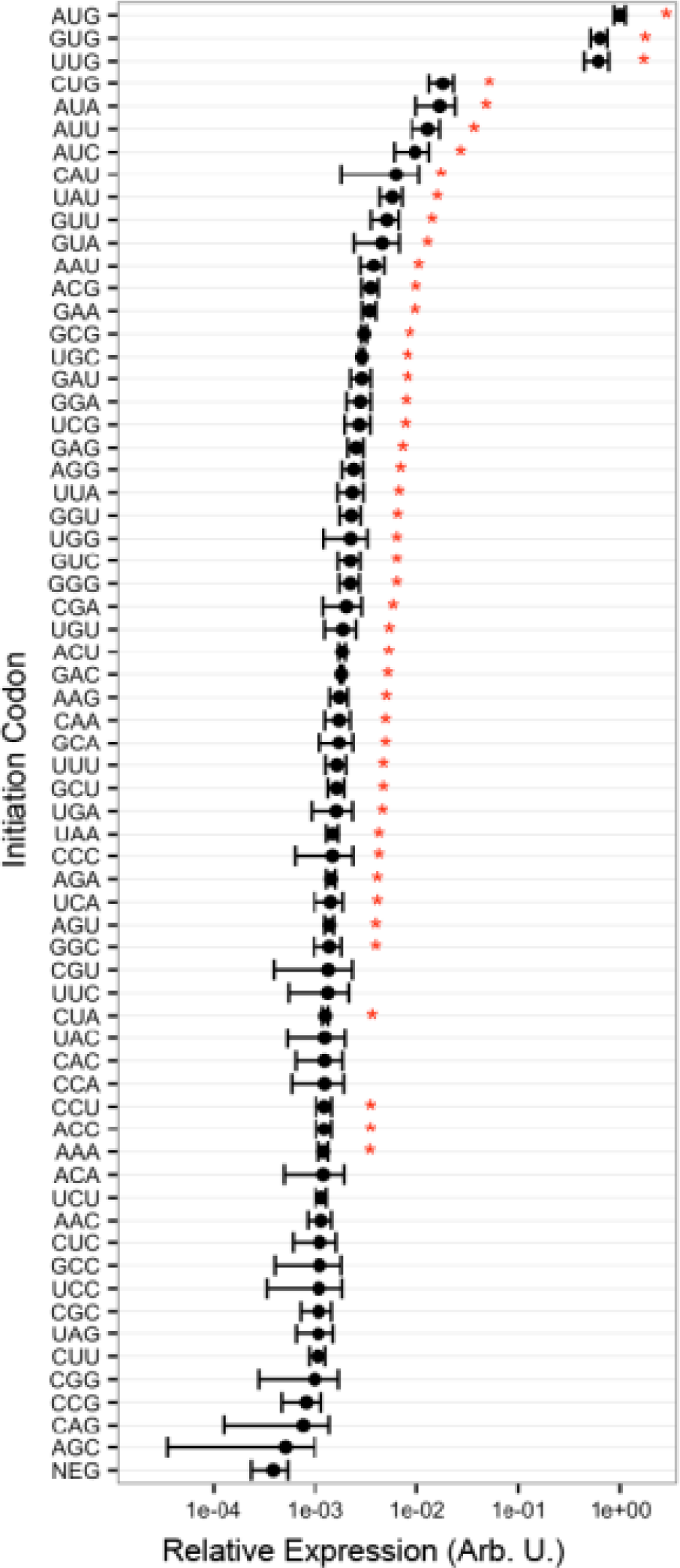
Translation initiation from all 64 codons. Mean per-cell fluorescence measured from cultures with each of the 64 codons as the start codon in the sfGFP coding sequence. Mean per-cell fluorescence of each culture was normalized by the mean per-cell fluorescence from the canonical AUG start codon. All points represent the mean and standard deviation of three biological replicates. Error bars represent one standard deviation. Red asterisks represent sfGFP expression significantly greater (adjusted p < 0.05) than the negative control (NEG) as determined by Dunnett’s test.

We wanted to confirm that the bulk fluorescence measurements we obtained on the plate reader were not arising from a small number of highly-expressing cells. We measured the distribution of fluorescence within the population of each culture on a flow cytometer. All cell populations were unimodally distributed (Supplementary Figure S3). Additionally, the geometric means of the fluorescence of these populations were well correlated with the mean per-cell fluorescence measured on the plate reader (Supplementary Figure S4).

Our initial experimental data suggest that the 64 codons can be organized into three groups: three canonical start codons (AUG, GUG, UUG), from which translation initiated at 10%-100% of AUG; four near-cognates (AUA, AUC, AUU and CUG), from which translation initiated at 1%-2% of AUG, similar to previously-reported values (52, 53); and 57 codons, from which translation initiated at 0.1 %-1% of AUG.

We examined the sfGFP coding sequence for in-frame upstream and downstream start codons as a possible explanation for the observed fluorescence. We found no canonical in-frame start codons upstream of the sfGFP coding sequence. There is an in-frame GUG at the 16th codon in the sfGFP coding sequence, but any resulting protein would be truncated and non-fluorescent because the minimal sequence needed for sfGFP fluorescence begins at the sixth codon (66).

We used proteolytic digest and mass spectrometry to determine if translation began at the modified start codons. Genes for sfGFP with AUC, ACG, CAU, GGA, and CGC (representing different levels of expression strength) as the start codons were re-cloned with a C-terminal 6x-His tag. After expression and purification, significant amounts of green protein were recovered from each culture except for the one with CGC as the start codon, which was also the only one of the five selected codons that did not initiate significant expression (Figure 2). We digested proteins with AspN and analyzed the mixture via mass spectrometry. Each expressed protein released peptides of intact N-termini, including an N-terminal methionine (Supplementary Tables S6-10). While ACG and AUC are one base away from AUG, CAU and GGA would require three point mutations to code for methionine. In one culture, with ACG as the start codon, there was also evidence of N-terminal peptides with the cognate amino acid, threonine (*M*r = 119), with a mass shift of −30 Da relative to methionine (*M*r = 149) (Supplementary Table S7). The mass spectrometry data suggest that the fluorescence we observed is from full-length GFP molecules expressed from non-AUG start codons with methionine as the N-terminal amino acid in the vast majority of synthesized proteins. Other researchers have also observed methionine in the N-terminal position of proteins whose translation initiates from GUG or UUG start codons (64, 67, 68).

We next explored translation initiated from non-canonical start codons under more physiologically relevant conditions. We focused on 12 codons spanning the observed expression initiation range (AUG, GUG, UUG, AUA, AUC, AUU, CUG, CAU, CGC, GGA, UAG and UGC). We cloned each of the 12 codons into first position of the coding sequence in three constructs: sfGFP under the control of the *RhaP_BAD_* rhamnose-inducible native *E. coli* promoter on a low-copy p15A plasmid (Figure 1B); NanoLuc luciferase under the control of the *RhaP_BAD_* promoter on a low-copy p15A plasmid (Figure 1C); and, NanoLuc luciferase on a single-copy bacterial artificial chromosome (Figure 1D). The same RBS and 5’-spacer was used in all constructs. sfGFP expression was quantified by measuring mean per-cell fluorescence. Luciferase expression was quantified by measuring mean per-cell luminescence emitted from the NanoLuc-catalyzed conversion of furimazine to furimamide (69). All measurements were repeated in triplicate. Measurements from serial dilutions of NanoLuc-expressing cells indicated that, over the range of concentrations used in this work, luminescence was linear with NanoLuc concentration (Supplementary Figure S5).

Our attempts to measure non-canonical sfGFP translation initiation driven by the *RhaP_BAD_* promoter on the low-copy p15A plasmid (Figure 1B) were impeded by a low signal-to-noise ratio due to significant background signal (Supplementary Figure S6). We were only able to detect significant sfGFP expression for the three canonical start codons AUG, GUG, and UUG (Figure 3A). We therefore transitioned to an expression system with lower background signal to detect non-canonical translation initiation under more biologically relevant expression conditions. NanoLuc luciferase was a good reporter for this application, because cell cultures emit negligible background luminescence, and the NanoLuc luciferase luminescence assay has a linear dynamic range greater than six orders of magnitude (69).

**Figure 3.**
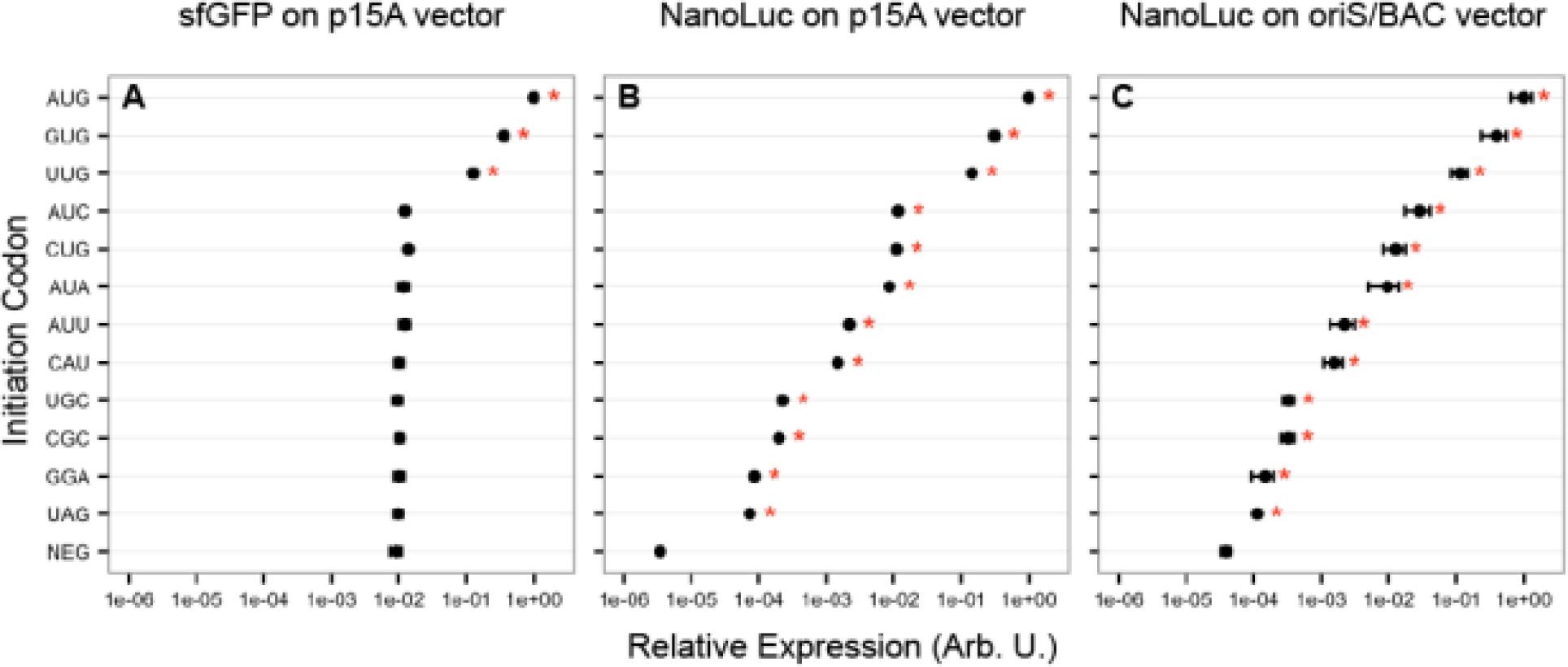
Translation inititation from a subset of 12 codons spanning the expression range. Translation initated from three expression cassettes, (A) sfGFP on a low-copy p15A plasmid, (B) NanoLuc luciferase on a low-copy p15A plasmid, and (C) NanoLuc luciferase on a single-copy bacterial artifical chromosome (BAC). All expression was driven by the *RhaP_BAD_* rhamnose-inducible native *E. coli* promoter. Per-cell (A) fluorescence and (B-C) luminescence was normalized by the expression from the canonical AUG start codon. All points represent the mean and standard deviation of three biological replicates. Error bars represent one standard deviation. NEG is the negative control cell. Red asterisks represent expression significantly greater (adjusted p < 0.05) than the negative control (NEG) as determined by Dunnett’s test.

We measured NanoLuc expression initiated from the 12 start codons listed above under transcriptional regulation of the *RhaP_BAD_* promoter on a low-copy p15A plasmid (Figure 1C). We were able to detect significant translation initiation from all 12 codons (Figure 3B). Translation initiated from the three canonical start codons at 10%-100% of AUG, from the four near-cognates at 0.1%–3% of AUG, and from the remaining five codons at 0.01%-0.1% of AUG. Of particular note was that translation initiated from UAG (a canonical stop codon) was the lowest of the 12 codons (0.007% of AUG), but was still more than an order of magnitude higher than the negative control (0.0003% of AUG).

We next measured NanoLuc expression from the single-copy bacterial artificial chromosome (BAC). As with the p15A plasmid (Figure 3B), we measured significant expression initiated from all of the 12 start codons (Figure 3C). As expected from the lower copy number, the absolute luminescence measured from constructs on the BAC was more than an order of magnitude lower than the absolute luminescence measured from the same constructs on the p15A plasmid. Translation initiated from the three canonical start codons at 10%-100% of AUG, from the four near-cognates at 0.02%–3% of AUG, and from the remaining five codons at 0.01%-0.2% of AUG. The lowest NanoLuc expression was again initiated from the canonical stop codon UAG (0.01% of AUG), which was still greater than the negative control (0.004% of AUG).

The relative strength of NanoLuc translation initiated from these 12 codons on the p15A plasmid, the relative strength of NanoLuc translation initiated from these 12 codons on the BAC, and sfGFP translation initiated on the pET20b plasmid were all well correlated (Supplementary Figure S7).

N-terminal RNA structure is known to impact translation initiation (70). We simulated RNA secondary structure around the initiation codon for all reporter plasmids used in this work (Figure 1) to evaluate if RNA secondary structure might contribute to translation initiation from non-canonical start codons. Both NUPACK (61) and KineFold (62) tools showed no correlation between the expected stability of the lowest energy structures and reporter expression for the NanoLuc constructs, and a weak correlation for the pET20b-sfGFP vector (Supplementary Figure S8 and Table S4). Additionally, there was no correlation between initiation codon GC-content and reporter expression (Supplementary Figure S8 and Table S4). These data suggest that differences in translation initiation from the start codons measured in this study were likely not caused by changes in RNA structure around the initiation codon or the GC-content of the initiation codon.

## DISCUSSION

In this work we have shown that translation initiates from many non-canonical start codons, both near-cognates and non-near-cognates, at levels ranging from 0.01–3% of translation initiated from the canonical AUG start codon in *E. coli.* The vast majority of these codons have never before been identified as start codons in *E. coli.* Given that average per-cell abundances of proteins in bacteria and mammalian cells span five to seven orders of magnitude (71, 72), translation initiation from non-canonical start codons may be more relevant than previously thought.

Past work in yeast has shown that there may not be sharp boundaries between genes and non-genic ORFs (73). We can imagine a scenario wherein, over evolutionary time scales, point mutations could create a weak non-canonical initiation codon downstream of a RBS. The small amounts of protein produced from this ORF, if beneficial to the organism, could select for further mutations that increased translation efficiency up to a point where the gene product more directly impacted organismal fitness. Further mutations could then be selected that tune for optimal expression dynamics in a given genetic context. Some evidence for this phenomenon exists in our start codon survey (Supplementary Table S5), in which the start codon of the *pcnB* (plasmid copy number B) gene alternate between AUG, GUG, and UUG across the bacterial phyla Cyanobacteria, Proteobacteria, Chlamydiales, and Hyperthermophiles. The *pcnB* gene encodes poly(A)polymerase, which is one of two redundant genes involved in adding polyadenine tails to the 3’ end of mRNA transcripts (74). The polyadenylation of an RNAi transcript regulates the copy number of the ColE1 plasmid origin used in this study (75).

One possible explanation for translation initiation from non-canonical start codons could involve non-specificity or conditional failure of regulatory mechanisms during translation initiation. Start codon selection is typically influenced by the proximity of the start codon to the ribosomal P-site, as determined by the binding of the 16S rRNA to the Shine-Dalgarno sequence (76). Binding of the tRNA^fMet^ anticodon to the start codon on the mRNA stabilizes the complex between the mRNA, tRNA^fMet^, IF2, and the 30S ribosomal subunit, allowing the 50S subunit to bind (17). IF3 acts as a gatekeeper in this process, allowing stabilization only when an acceptable start codon is bound (52). Mutations in both IF3 and the 16S rRNA can increase translation initiation from non-AUG start codons (53, 77, 78).

Our data reaffirm AUG, GUG and UUG as the strongest start codons in *E. coli.* However, we wonder why so few non-canonical codons have been annotated in bacterial genomes. One possibility is that naturally occurring non-canonical start codons are in fact exceedingly rare. Another possibility is that many naturally occurring non-canonical start codons and so-initiated proteins remain undiscovered. In a recent *E. coli* whole cell shotgun proteomics experiment, approximately half of the detected spectra could not be mapped to known genes (79). The presence of frequent but very low-level expression of proteins via non-canonical start codons would have widespread implications for genome annotation, cellular engineering, and our fundamental understanding of translation initiation.

## ACKNOWLEDGMENTS

The authors would like to acknowledge Atri Choski, Geremy Clair, Steven Hallam, Sarah Munro, and Ljiljana Pasa-Tolic, for helpful discussions, and Christine Chang for experimental assistance. The authors would like to thank Steve Lund for assistance with statistical data analysis. Cell sorting/flow cytometry analysis for this project was done on instruments in the Stanford Shared FACS Facility, with particularly helpful assistance from Marty Bigos and Cathy Crumpton. We would like to acknowledge Ryan Leib and Chris Adams at the Vincent Coates Foundation Mass Spectrometry Laboratory, Stanford University Mass Spectrometry (http://mass-spec.stanford.edu) for assistance in protein analysis. The authors would like to acknowledge Sara Lefort and the Friday morning Coffee Hour sponsored by the Ginzton Lab at Stanford University for providing the venue that facilitated the key conversation that inspired this project. The bacterial artificial chromosome was a generous gift from Fernan Federici of the Universidad Catolica de Chile. The authors acknowledge the financial support of the NRC/NIST Postdoctoral Research Program.

## Footnotes

i Certain commercial equipment, instruments, or materials are identified in this report to specify adequately the experimental procedure. Such identification does not imply recommendation or endorsement by the National Institute of Standards and Technology, nor does it imply that the materials or equipment identified are necessarily the best available for the purpose.

